# Moving toward versus away from another: how body motion direction changes the representation of bodies and actions in the visual cortex

**DOI:** 10.1101/2020.08.06.239749

**Authors:** Emmanuelle Bellot, Etienne Abassi, Liuba Papeo

## Abstract

Representing multiple agents and their mutual relations is a prerequisite to understand social events. Using functional MRI on human adults, we show that visual areas dedicated to body-form and body-motion perception contribute to processing social events, by holding the representation of multiple moving bodies and encoding the spatial relations between them. In particular, seeing animations of human bodies facing and moving toward (*vs*. away from) each other, increased neural activity in the body-selective cortex (extrastriate body area -EBA) and posterior superior temporal sulcus for biological-motion perception (bm-pSTS). In those areas, representation of body postures and movements, as well as of the overall scene, was more accurate for facing-body (*vs*. non-facing body) stimuli. Effective connectivity analysis with Dynamic Causal Modeling revealed increased coupling between EBA and bm-pSTS during perception of facing-body stimuli. The attunement of human vision to multiple-body scenes involving perceptual cues of interaction such as face-to-face positioning and approaching behaviour, was further supported by the participants’ better performance in a match-to-sample task with facing-body *vs*. non-facing body stimuli. Thus, visuo-spatial cues of interaction in multiple-person scenarios affect the perceptual representation of body and body motion and, by promoting functional integration, streamline the process from body perception to action representation.

## INTRODUCTION

How does the brain construct the representation of events? Even in the simple case of two bodies looking at each other, the meaning of an event is not as much in the appearance or the identity of the two, as in the relation that connects, binds and assigns them to a role, to determine *who does what to whom*.

In the visual world, the presence of certain visuo-spatial relations in a scene may contribute to channel object/scene perception into event representation. Research on the visual perception of complex scenes involving multiple bodies (i.e., people) has shown that the same scene is processed differently, depending on the spatial relation between bodies in the scene: visual perception is particularly efficient for scenes featuring relations that recur in social interactions, such as spatial proximity and face-to-face positioning of bodies. Indeed, relative to unrelated bodies, seemingly interacting bodies have privileged access to visual awareness – i.e., they are more likely to be detected and recognized– when presented with visual noise (Papeo et al. 2017a), or in a crowded environment (Papeo et al. 2019; Vestner et al. 2019). Moreover, spatial relations cuing interaction facilitate the extraction of information about action (Neri et al. 2006; Glanemann et al. 2016), and participants in the action (e.g., agent/patient roles; Hafri et al. 2013, 2018).

In line with those studies, functional MRI (fMRI) research has shown that the body-selective visual cortex responds to nearby face-to-face bodies more strongly than to the same bodies presented back-to-back; and it discriminates body postures better in a face-to-face configuration than in a back-to-back configuration (Abassi and Papeo 2020). Among all the visual areas, those effects were the most reliable in the extrastriate body area (EBA), a region in the lateral occipital cortex, specialized to perception of bodies and body parts (Downing et al. 2001; Peelen and Downing 2005). Overall, those results suggested that a body is represented in different ways depending on whether, or not, it faces toward another and is reciprocated; if it does, its representation in the visual cortex is enhanced.

The EBA response to facing bodies may reflect a key mechanism, whereby visual perception contributes to the formation of social-event representation by capturing and emphasizing visuo-spatial cues of interaction in multiple-person scenes. In the vast majority of cases, social interaction involves bodies and body parts in movement. Therefore, the mechanism identified in the EBA could drag up to the visual cortex specialized to perception of body motion and action.

Neuronal populations in the posterior superior temporal sulcus (pSTS) consistently responds to visual biological motion more strongly than to mechanical motion (Beauchamp et al. 2003; Pyles et al. 2007; Grosbras et al. 2012), and to human-like bipedal motion more strongly than to quadrupedal motion (Papeo et al. 2017b), with discrimination of body motion direction (e.g., forward *vs*. backward; Vangeneugden et al. 2014). Disruption of activity in the biological-motion posterior superior temporal sulcus (bm-pSTS) impairs the ability to reconstruct action from motion patterns (Grossman et al. 2005; Saygin 2007). Thus, integrating signals from category-specific regions like the EBA, the bm-pSTS critically contributes to forming a representation of human motion and action (Grossman and Blake 2002; Vangeneugden et al. 2014). For its functional properties, the bm-pSTS is also considered the main entry into the social brain network (Allison et al. 2000; Hein and Knight 2008; Graziano and Kastner 2011; Lahnakoski et al. 2012), a view further promoted by the proximity to areas implicated in social cognitive functions such as theory of mind (Saxe and Powell 2006; Saxe et al. 2006), and in the representation of social-interaction contents conveyed by movies of acting people, point-light displays of human-body dyads, as well as animated geometrical shapes (Pelphrey et al. 2004; Sinke et al. 2010; Centelles et al. 2011; Georgescu et al. 2014; Van den Stock et al. 2015; Isik et al. 2017; Walbrin et al. 2018; Walbrin and Koldewyn 2019).

The extent to which the bm-pSTS is implicated in forming a representation of social events, and how it would do so, remains unknown. It is unknown whether this region can hold the representation of multiple moving bodies and, if so, whether it cares about relations between different bodies and movements performed by different bodies, before information is sent forward for integrative processes toward action understanding.

Here, we asked how brain areas, key for visual perception of body form and body motion, encode dynamic scenes featuring multiple bodies acting and moving toward one another *as if* interacting (facing dyad), or acting and moving away from one another (non-facing dyads). In both facing and non-facing dyads, the two bodies performed different movements/actions simultaneously, but they did not touch or interact in any meaningful way. This methodological choice allowed testing the specific hypothesis behind the study, according to which visual perception areas encode for visuo-spatial cues associated with social interaction (i.e., spatial proximity, body orientation and approaching behaviour), rather than the social interaction itself. Based on the above results (Papeo et al. 2017a; Papeo and Abassi 2019; Vestner et al. 2019; Abassi and Papeo 2020), we expected to find an advantage in processing facing dyads, which we sought to capture in the neural activity (through univariate and multivariate analyses of the fMRI signal, Study 1), as well as in the participants’ performance in visual recognition of multiple-body scenarios (Study 2).

Finally, we studied whether visuo-spatial cues of interaction in multiple-body scenarios (i.e., facing *vs*. non-facing positioning) affected the coupling between the EBA and the bm-pSTS. Coupling was determined by effective connectivity estimated with dynamic causal modelling (DCM), which informs on how two regions influence each other during stimulus processing (Friston et al. 2003). Increased coupling between the EBA and the bm-pSTS in the facing-dyad (*vs*. non-facing) condition would contribute to demonstrating that, in a multiple-body scene, visuo-spatial cues of interaction not only affect the perceptual representation of bodies and body motion in the dedicated brain areas but, by promoting functional integration, they also streamline the process from body perception to action representation.

## MATERIALS AND METHODS

### Study 1: fMRI

#### Participants

Twenty healthy adults (11 females, mean age 25 ± 4 years) participated in the fMRI study as paid volunteers. All participants had normal or corrected-to-normal vision and reported no history of medical, psychiatric or neurological disorders, or use of psychoactive medications. They were screened for contraindications to MRI and gave informed consent before participation. The study was approved by the local ethics committee (Comité de Protection des Personnes Sud Est V, CHU de Grenoble).

#### Stimuli

Two-second (s) silent movies of point-light displays were created from the “Communicative Interaction Database - 5AFC format” (CID-5; Manera et al. 2016). The database includes movies featuring an interaction between two human bodies, each defined by 13 white dots in correspondence with the major reference points (top of the head, shoulders, elbows, hands, hips, knees and ankles). From those movies, we isolated 28 single bodies performing an action with an invisible object or without an object, and asked a group of participants external to the fMRI experiment (n=30), to describe the action of each single body presented alone. Ambiguous stimuli, i.e. those whose description did not match the description in the original database, were eliminated. The twenty bodies that obtained the most consistent description across participants were selected to create ten dyads in which two bodies acted simultaneously, with no contact or obvious meaningful interaction. Based on the selected bodies the following three sets were created: 1) ten facing dyads, in which the two bodies moved and acted toward each other; 2) ten non-facing dyads, obtained by horizontally rotating of 180 degree, each body in the facing dyads, so that the two moved and acted away from each other; 3) 20 single bodies that formed the dyads, facing and moving leftward (50%) or rightward.

While bodies moved toward each other in facing dyads and away from each other in non-facing dyads, we ascertained that the average distance between the two bodies was matched across the two conditions. To verify so, for each movie, we selected the frames in which the two bodies were the farthest (F1) and the closest (F2) (see Figure 1). In each selected frame, the eight most informative points of each body (i.e., shoulders, hands, hips, ankles) were used to compute the polygon centroid point of the body. The distance between the two bodies in a movie was quantified as the number of pixels along the x-axis (horizontal distance) between the centroids of the two bodies at F1 and F2 (i.e., D1 for F1 and D2 for F2): D = |D1 − D2|. Statistical analysis (two-tailed *t*-test) showed that distance D between the two bodies was comparable for facing and non-facing dyads (mean_facing_ = 85 pixels ± 82 SD; mean_non-facing_ = 92 pixels ± 79 SD; *t*(9) < 1, *ns*). In summary, since identical bodies and body movements were presented, with matched distances across conditions, the relative spatial positioning of bodies was the only feature that consistently differed between facing and non-facing dyads.

**Figure 1.**
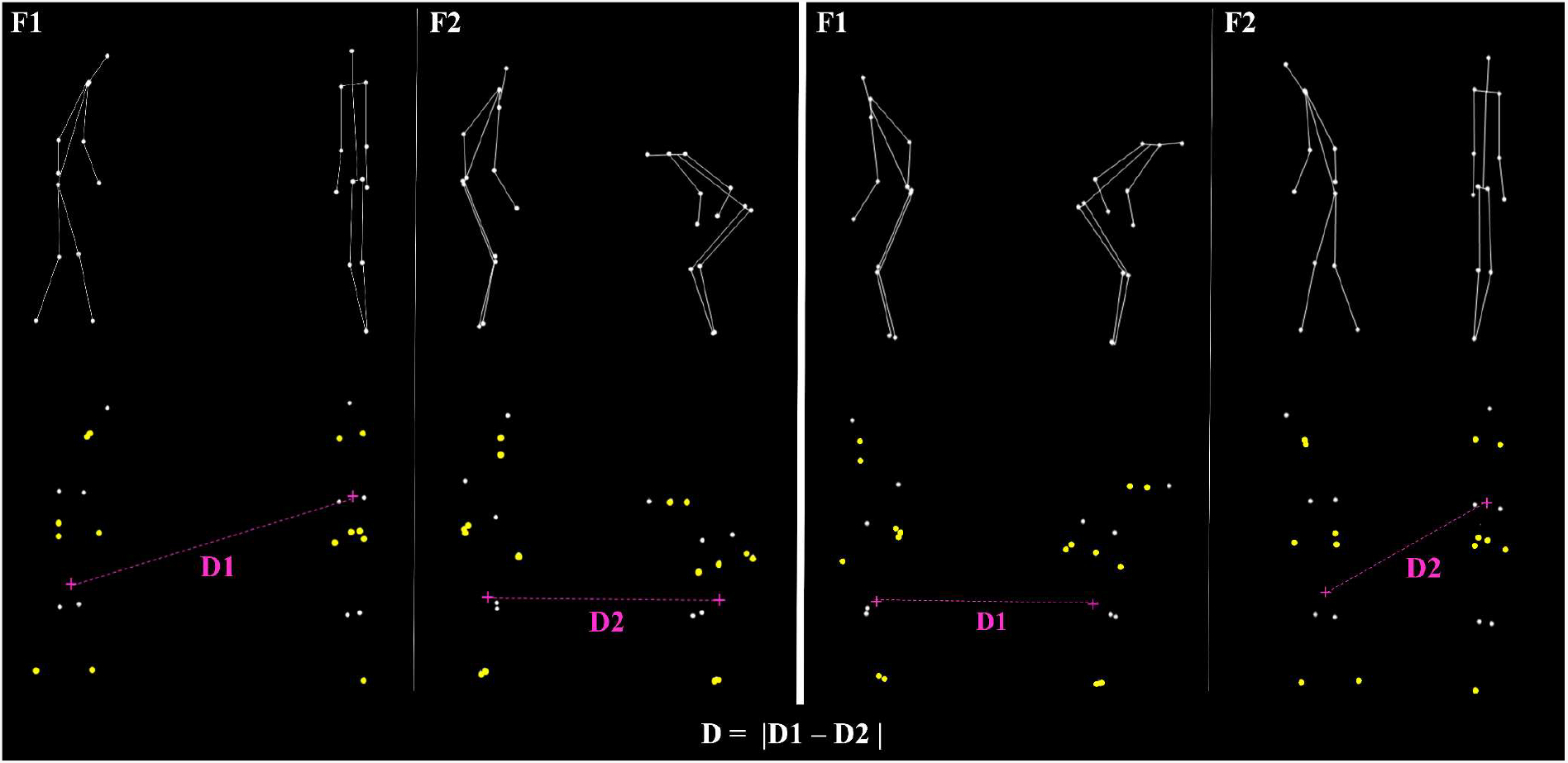
Example of point-light displays used in Study 1-2 (dots are connected by full lines for illustration; dots used to measure the distance between two bodies are highlighted in yellow). *Top*. Example of frames selected from a facing-dyad video-clip (left) and the corresponding non-facing dyad video-clip (right), to compute the distance between the two bodies: F1 is the frame in which the two bodies are the farthest; F2 is the frame in which the two bodies are the closest. *Bottom*. Highlighted in yellow are the eight points for each body used to compute the polygon centroid point, indicated by a pink cross. For F1 and F2, distance (quantified as the number of pixels along the x-axis) between the centroids of the two bodies (D1 for F1 and D2 For F2) was measured. The distance value for a given movie was defined as the absolute difference (D) between D1 and D2.

All the stimuli were horizontally flipped to create new stimuli, yielding a total of 80 stimuli (40 original stimuli and their flipped version), so distributed across conditions: 20 facing dyads, 20 non-facing dyads, 20 rightward single bodies, 20 leftward single bodies. All the stimuli were edited as .avi files with a resolution of 640 × 480 pixels and a frame rate of 30 frames/sec.

#### Experimental design and procedure of the fMRI study

The fMRI study included two parts, a main experiment and three functional localizer tasks, all performed during fMRI.

##### Main fMRI experiment

The experiment consisted of an event-related design involving three conditions: single bodies, facing dyads and non-facing dyads, all presented in the form of point-light displays (see Stimuli). Facing and non-facing dyads were randomly presented over two functional runs, each lasting 8 minutes (min) and 32 s. Each run consisted of two main sequences of visual stimulation separated by an interval of 16 s. Each sequence included forty 2-s events (movies of facing and non-facing dyads) with variable inter-stimulus interval (ISI) of 2, 4 or 6 s, each occurring with 1/3 probability. Events in the first sequence were presented in random order, and events in the second sequence were presented in the counterbalanced (i.e., reversed) order relative to the first sequence. Thus, events that were presented at the end of the first sequence were shown at the beginning of the second sequence, and *vice versa*. Each sequence featured 2 repetitions of the same stimulus (original view and flipped version). Therefore, each movie-stimulus was presented 4 times in each run for a total of 8 times across the whole experiment. The first sequence of a run ended with 16 s of fixation (fixation cross at the center of the screen), which allowed splitting each run in two independent chunks for the multivariate analyses (see below). Each run began with a warm-up block (8 s) and ended with a cool-down block (16 s), during which a central fixation cross was presented. The experiment included two additional runs identical to the above two runs, except that, instead of facing and non-facing dyads, leftward and rightward single bodies stimuli were shown at the center of the screen.

In a subset of events across a run (2.5%), dots forming the point-light displays changed colour (from white to light pink). Participants were instructed to fixate the center of the screen and to report the colour change as soon as it occurred, by pressing a button on a remote placed under the right index finger. Those task instructions intended to minimize eye movements and maintain vigilance in the scanner. The main experiment including two runs of dyads and two runs of single bodies lasted 34 min and 8 s.

Stimuli in this task and in the following functional localizer tasks were back-projected using an LCD projector (frame rate: 60 Hz, screen resolution: 1024 × 768 pixels, screen size: 40 × 30 cm) onto a translucent screen positioned at the rear of the magnet. Participants viewed the screen binocularly (7° of visual angle) through a mirror above the head coil. Stimulus presentation, response collection and synchronization with the scanner were controlled with the Psychotoolbox (Brainard 1997; Pelli 1997) through MATLAB (MathWorks, MA, USA).

##### Functional localizer tasks

After the main experiment, we ran two functional localizer tasks (Lingnau and Petris 2013; Papeo and Lingnau 2015) to define the bm-pSTS region that responds to biological motion, and the middle temporal visual area (MT/V5) that responds to motion, respectively. Moreover, 15 of the 20 participants included in the study took part in a second fMRI session in which they performed a functional localizer task to identify the EBA.

###### Functional localizer task for the bm-pSTS

Participants watched blocks of white point-light displays on a black background, depicting human body movements (6 blocks) or abstract animations (6 blocks) obtained by scrambling the global configuration of dots that formed the human body movements, while keeping identical the local motion of each dot. Each point-light display lasted 1.5 s and was followed by 0.5 s of ISI. Blocks lasted 14 s and were separated by 14 s of fixation. Blocks of body movements and abstract animations were randomly interleaved in one single run. The run began with a 14-s warm-up block and ended with a 14-s cool-down block, during which a blank was shown. The task lasted 5 min and 50 s.

###### Functional localizer task for the MT

The design of this task was identical to the above functional localizer task. In this task, point-light displays showed moving or static white dots on a black background. Moving dots moved outwards along the radial axis at a speed of 4 deg/s. This task was performed for purposes external to the current study and won’t be discussed further.

###### Functional localizer task for the EBA

This task was adapted from the fLoc package (Stigliani et al. 2015) developed to identify category-specific activity in the visual cortex. In this task, participants viewed 180 grayscale photographs of five object classes (whole bodies and body parts, faces, places, inanimate objects and scrambled objects). Stimuli were presented over two runs of 5 min and 16 s each. A run included 72 blocks of 4 s each, 12 for each object class (bodies, faces, places, inanimate and scrambled objects) with eight images per block (500 ms per image without interruption), randomly interleaved with 12 baseline blocks showing a blank. Each run began with a 12-s warm-up block and ended with a 16-s cool-down block. Participants were instructed to press a button when they saw two identical stimuli separated by another stimulus (2-back repetition detection task).

##### Data acquisition

Experiments were performed using a whole-body 3-Tesla Magnetom Prisma MRI scanner (Siemens AG, Heathcare). For functional scans, T2*-weighted volumes were acquired using a multi-band accelerated echo planar imaging (EPI) pulse sequence (http://www.cmrr.umn.edu/multiband; repetition time 2 s, echo time 30 ms, 56 slices, slice thickness 2.2 mm, no gap, field-of-view 210 × 201 mm, flip angle 80°, acquisition matrix 96 × 92, multiband acceleration factor of 2 with phase encoding set to anterior/posterior direction). High-resolution T1-weighted anatomical images were collected using a MPRAGE pulse sequence (224 sagittal slices, repetition time 3 s, echo time 3.7 ms, inversion time 1.1 s, flip angle 8°, acquisition matrix 320 × 280, field-of-view 256 × 224 mm, slice thickness 0.8 mm, GRAPPA accelerator factor of 2). For all the participants, the fMRI session had the following organization: two functional runs of the main experiment (256 images each), acquisition of anatomical images (8 min), two functional runs of the main experiment (256 images each), one run of the functional localizer task for the pSTS (175 images) and one run of the functional localizer task for the MT (175 images). The functional localizer task to define the EBA was performed on a different day, before or after the above session, with the same parameters for acquisition as above. Framewise Integrated Real-time MRI Monitoring (FIRMM; Dosenbach et al. 2017) was used during MRI, to monitor the data quality in real time. During the acquisition, the participant laid supine in the scanner, with the head surrounded by soft foam to reduce head movements. The cardiac signal was indirectly recorded based on the hemodynamic pulse at fingertip using the pulse plethysmography unit attached to the MRI scanner and used as covariate of non-interest.

#### Data analyses

##### Pre-processing

The first four volumes of each run were discarded, taking into account initial scanner gradient stabilization (Soares et al. 2016). Pre-processing of the remaining volumes involved geometric distortion correction (using fieldmaps), slice time correction, spatial realignment and motion correction using the first volume of each run as reference. The maximum displacement was 0.54 mm on the x-axis (mean_max_ = 0.23, SD = 0.13), 2.16 mm on the y-axis (mean_max_ = 0.56, SD = 0.49) and 2.20 mm on the z-axis (mean_max_ = 0.85, SD = 0.51). Anatomical volumes were co-registered to the mean functional image, segmented into grey matter, white matter and CSF in native space, and normalized in the Montreal Neurological Institute (MNI) space. A default fourth degree B-spline interpolation was applied. The parameters for anatomical normalization were used for the normalization of functional volumes. Finally, each functional volume was smoothed by a 6-mm FWMH (Full Width at Half Maximum) Gaussian kernel for univariate analysis and by a 2-mm FWMH Gaussian kernel for multivariate analysis. Times-series for each voxel were high-pass filtered (1/128 Hz cutoff) to remove low-frequency noise and signal drift.

##### Univariate analyses

###### Whole-brain analysis

Statistical analysis of the pre-processed blood-oxygen-level-dependent (BOLD) signal was performed using a General Linear Model (GLM) (Friston et al. 1994) in SPM12 (Friston 2007) and MATLAB (MathWorks, MA, USA). Conditions of interest (single bodies, facing dyads, non-facing dyads and fixation periods) were modelled as regressors, constructed as boxcar functions convolved with the canonical hemodynamic response function (HRF). Subject’s responses to the colour-change detection task (correct responses or errors) and the movement parameters derived from realignment corrections (three translations and three rotations) were entered in the design matrix as additional factors of no interest. To identify the brain networks that responded to facing and non-facing body dyads, statistical images corresponding to the contrasts [Facing > Non-facing dyads] and [Non-facing > Facing dyads] were computed for each participant. To account for inter-subject variability, contrast images of all the participants (level 1) were included in a second level *t*-test (voxelwise threshold *p* < 0.001).

###### Regions of interest (ROIs) definition

Using the data from the functional localizer tasks, we identified in each participant two ROIs: the bm-pSTS area sensitive to visual biological motion and the EBA sensitive to visual body shapes. As controls for the specificity of the effects in the target ROIs, we identified for each individual two additional ROIs, the early visual cortex (EVC) and the parahippocampal place area (PPA).

###### bm-pSTS

For each participant, the bm-pSTS was defined by entering the data registered during the functional localizer task into a GLM with two regressors of interest (Intact point-light, Scrambled point-light) and six regressors for movement correction parameters as nuisance covariates. The left and right bm-pSTS were identified with the contrast Intact > Scrambled point-light displays. All the voxels that passed a threshold of *p* = 0.05 were ranked by activation level based on *t* values. The final left and right ROIs included the 200 best voxels.

###### EBA

For 15 of our participants, the EBA was defined by entering the individual data registered during the functional localizer task into a GLM with two regressors of interest (bodies, objects), one regressor for baseline blocks and six regressors for movement correction parameters as nuisance covariate. Bilateral masks of the inferior lateral occipital cortex (LOC) were created using FSLeyes (McCarthy 2019) and the Harvard-Oxford Atlas (Desikan et al. 2006) through FSL. (Jenkinson et al. 2012). Within these masks, left and right EBA were identified with the contrast Bodies > Objects. All the voxels that passed a threshold of *p* = 0.05 were ranked by activation level based on *t* values. The final left and right ROIs included the 200 best voxels. For the remaining participants who did not perform this task (n = 5), EBA was defined by selecting, for the contrast of interest, the 200 best voxels obtained at the group level (second level).

###### PPA

Using the data collected during the functional localizer task for the EBA, we defined the PPA as a high-level control ROI, to assess the specificity of the effects of body positioning in the two target ROIs (EBA and bm-pSTS). PPA was defined in 15 participants, by entering the individuals’ data into a GLM with two regressors of interest (places, objects), one regressor for baseline blocks and six regressors for movement correction parameters as nuisance covariate. Bilateral masks of the inferior parahippocampal cortex were created using FSLeyes (McCarthy 2019) and the Harvard-Oxford Atlas (Desikan et al. 2006) through FSL (Jenkinson et al. 2012). Within these masks, left and right PPA were identified with the contrast Places > Objects. All the voxels that passed the threshold of *p* = 0.05 were ranked by activation level based on *t* values. Left and right ROIs included the 200 best voxels. For the participants who did not perform this task (n = 5), PPA was defined by selecting, for the contrast of interest, the 200 best voxels obtain at the group level (second level).

###### EVC

We defined the EVC as an additional control ROI. A bilateral mask of the EVC was created using a probabilistic map of visual topography in the human cortex (Wang et al. 2015). After transforming the mask in each subject’s space, the 200 voxels with the highest probability were selected to define the individual EVC-ROI.

###### ROIs analysis

Mean activity values (mean β-weights minus baseline) for facing and non-facing dyads were extracted separately from the left and right ROIs (bm-pSTS, EBA, PPA and EVC). These data were analysed in a 2 Hemisphere (left *vs*. right) × 4 ROI × 2 Spatial relation (facing *vs*. non-facing) ANOVA. Post-hoc comparisons between conditions were performed using pairwise *t*-tests (α = 0.05, two-tailed). All statistical analyses were performed with Statistica (StatSoft, Europe, Hamburg) and power analyses were realized using G*Power (Faul et al. 2009).

##### Multivariate pattern analyses (MVPA)

We used MVPA to examine, in the ROIs, the neural representation of dyads, as well as of single bodies, as a function of the spatial relation between the two bodies (facing *vs*. non-facing). To this end, in each ROI, we tested the discrimination of facing dyads and of non-facing dyads, and the discrimination of single-body movements seen in facing *vs*. non-facing dyads. We were especially interested in assessing whether visuo-spatial cues of interaction enhanced the perceptual representation of body dyads as wholes, as well as the representation of the individual body postures (in the EBA) and movements (in the bm-pSTS).

###### Multi-class classifications of facing and non-facing dyads

In each ROI, we tested the discrimination of classes (i.e., exemplars) of the same category of stimuli (facing or non-facing dyad). A β-pattern was estimated for each presentation of a given dyad; therefore, for each ROI and each participant, we obtained 80 β-patterns for facing dyads, 80 for non-facing dyads, and 160 for single bodies. β-patterns were normalized run-wise to avoid spurious correlations within runs (Lee and Kable 2018). For each single participant and each ROI, a support vector machine (SVM) classifier (LIBSVM, Chang and Lin 2011) implemented in the COSMOMVPA toolbox (Oosterhof et al. 2016) was trained to discriminate between the patterns associated with 10 classes of stimuli (i.e., 10 unique facing dyads classes), using 8 samples (i.e., patterns) per class. At each iteration, the classifier was trained on 3 out 4 chunks, and tested on the held-out chunk. For each participant and each ROI, we cycled through 4 held-out iterations and averaged the classification accuracy across all the iterations. Participants’ classification accuracies were entered in a one-sample *t*-test (α = 0.05, two-tailed), separately for each ROI, to assess whether the average classification accuracy was significantly better than chance. Because classification could be correct in one out of ten classes, the chance level was 10%. Identical training-and-test scheme was repeated for all ROIs, using the β-patterns measured for non-facing dyads. Classification accuracies for facing *vs*. non-facing dyads were compared using pairwise *t*-test (α = 0.05, two-tailed).

###### Multi-class classifications of single bodies from dyads

In this analysis, we asked how well a single body action could be discriminated when seen in a facing dyad or in a non-facing dyad. An SVM classifier was trained with the patterns of 20 different classes corresponding to the 20 single bodies, using eight samples per class. Using a one-against-one approach and voting strategy as implemented in LIBSVM (Chang and Lin 2011), the classifier was tested on the classification of the 80 patterns corresponding to facing dyads (8 samples for each of the 10 unique facing dyads). In 80 testing iterations, a pattern representing a facing dyad could be classified in two of the 20 classes of single bodies. Because the classification of each dyad could be correct in two out of 20 cases, the chance-level was 10%. For each ROI and each participant, one classification accuracy value was obtained by averaging across all the iterations. Participants’ accuracy values were entered in a one-sample *t*-test (α = 0.05, two-tailed), separately for each ROI, and evaluated against chance-level classification. The same analysis was repeated for all ROIs, using the patterns associated with non-facing dyads as the test set. Classification accuracies for facing *vs*. non-facing dyads were compared using pairwise *t*-test (α = 0.05, two-tailed).

##### Dynamic Causal Modeling analysis

Next, we asked whether visual perception of body dyads in the facing *vs*. non-facing configuration differentially modulated the connectivity that is established between the EBA and the bm-pSTS during body perception. We used Dynamic Causal Modeling (DCM) analysis to evaluate effective connectivity between the two functionally-defined ROIs, based on the fMRI time-series (Friston et al. 2003; Penny et al. 2004). This approach allows estimating how one neural system influences another and how such relationship is affected by the experimental conditions.

###### Time-series extraction

We studied how, during perception of dyads, the relationship between the bm-pSTS and the EBA changed as a function of the spatial relation between bodies in the dyads: facing *vs*. non-facing dyads. Time-series from the two ROIs were concatenated across the four runs at the individual level to obtain a single time-series including both the runs with single bodies and with dyads. For this analysis, ROIs were defined with the contrast [Visual stimulation > Fixation], whereby visual stimulation included single-body, facing dyad and non-facing dyad events. Activations were selected within the functional masks defined with the functional localizer tasks for the EBA and the bm-pSTS. For each participant and each ROI, a sphere of 6mm radius was defined around the activity peak within the mask. Subject-specific maxima inside these spheres were localized, and the principle eigenvariates (time series) adjusted for the participant’s effect of interest (i.e. single-body, facing dyad, non-facing dyad and fixation) were extracted.

###### Construction of model space

A base model composed of two hubs (bm-pSTS and EBA) was defined. Alternative models were derived from the base model. For each model, three parameters, all expressed in Hertz (Hz), were estimated: i) *driving input parameters*, reflecting how the ROIs respond to external stimuli; ii) *endogenous connection parameters*, reflecting the baseline effective connectivity between the ROIs (i.e., how the rate of activity changes in one target-region is influenced by the increase of activity in a source-region); and iii) *modulatory parameters*, quantifying how effective connectivity is influenced by the experimental context. Modulatory connections, obtained by adding modulatory parameters to endogenous parameters, reflect context-induced change in connectivity between source- and target-region. Positive parameter values indicate that the rate of activity change in the target-region increases, as the activity in the source-region increases; negative parameter values indicate that the rate of activity change in the target-region increases, as the activity in the source-region decreases.

We defined three models. Across the three models, the single-body condition was used as the driving visual input (external visual stimulation) that entered the system through the EBA. Endogenous connections between the EBA and the bm-pSTS could be unidirectional (EBA→bm-pSTS) or bidirectional (EBA↔bm-pSTS). Finally, we estimated the modulation of the unidirectional connection EBA→bm-pSTS and of the bidirectional connections EBA↔bm-pSTS as a function of the context: facing *vs*. non-facing. The three models were used to assess how the perception of facing and non-facing dyads modulated the endogenous connectivity between the EBA and the bm-pSTS, underlying body perception (see Figure 2). Two model spaces were created, both including the above three models but differing with respect to the type of modulation: modulation of endogenous connectivity by perception of facing dyads for the “Facing dyads” model space, and modulation of endogenous connectivity by perception of non-facing dyads for the “Non-facing dyads” model space.

**Figure 2.**
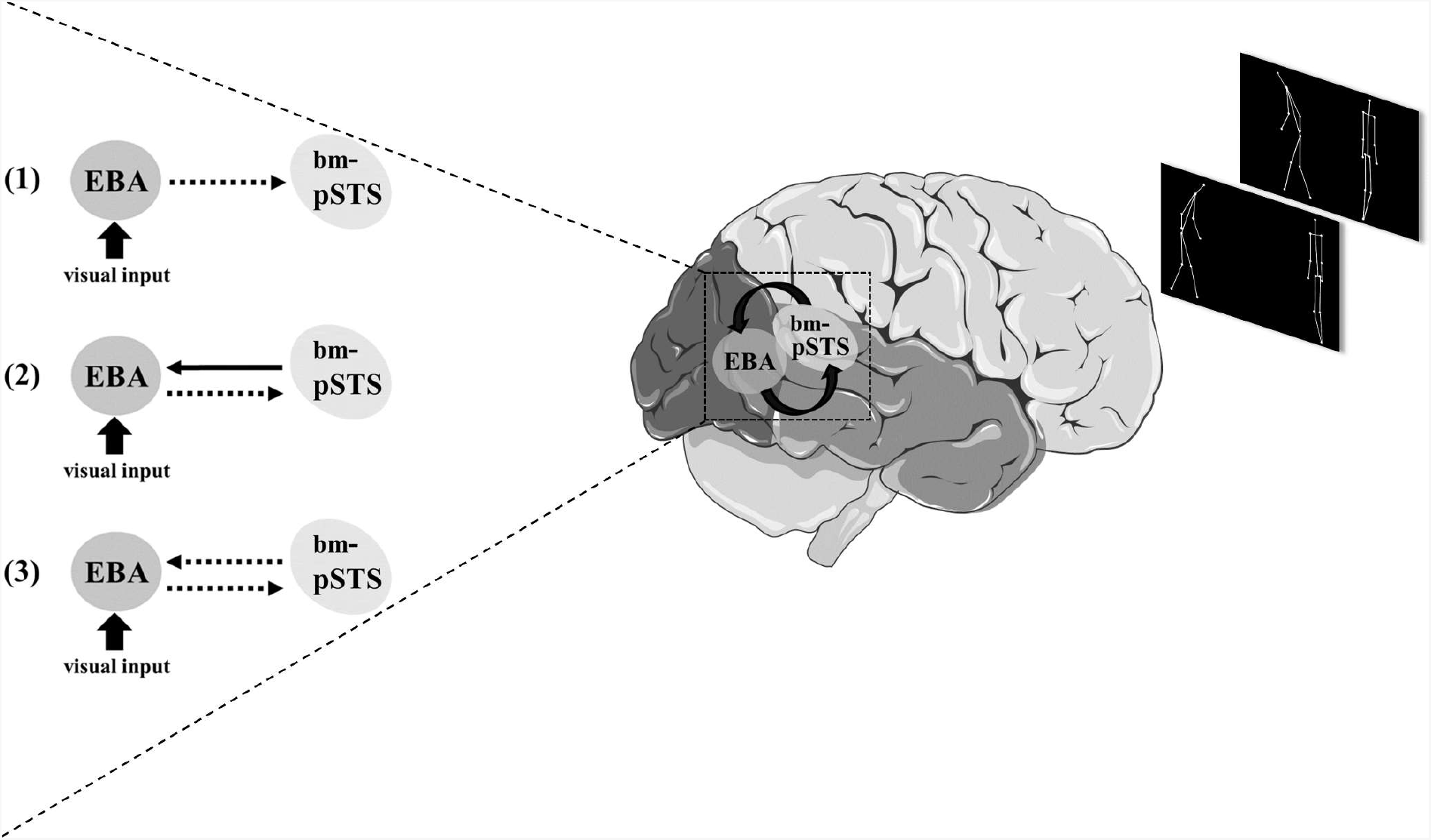
Illustration of the model space defined to study how the connectivity between the EBA and bm-pSTS changed as a function of the two experimental conditions: perception of facing *vs*. non-facing dyads. Two model spaces were created, “Facing dyads” and “Non-facing dyads” model spaces, both including the same three models (1 to 3), but differing for the type of modulation (modulation by facing dyad or by non-facing dyad perception). In each of the three models, the driving visual input (single bodies) is indicated by the thicker solid arrow. All other arrows represent endogenous connectivity. Dotted arrows indicate endogenous connections that could be modulated by (facing/non-facing) dyad perception. Model 1 involves a unidirectional connection from the EBA to the bm-pSTS, modulated by dyad perception; model 2 involves bidirectional connections between the EBA and the bm-pSTS with a modulation of the EBA to the bm-pSTS connection by dyad perception; and model 3 involves bidirectional connections between the EBA and the bm-pSTS both modulated by dyad perception.

###### Model estimation and comparison

For each participant, for each model, we computed the model evidence, which reflects the probability of obtaining the current neural data given a model. Model evidence was used to compare the different models and identify the one that best predicted the actual data. Random-effects Bayesian Model Selection (BMS) in the DCM tool of SPM12 (version 12.5) was used to calculate the probability of each model to be more likely than any other tested model at the group level (i.e., exceedance probability, xp) (Stephan et al. 2009). The model with the highest xp is considered as the best model for explaining the data. Generally, a model is selected when its xp exceeds 0.9 (i.e., the model is 90% more likely to explain the data than any other tested model). Since all xp values were below 0.9 (see Results), we used Bayesian Model Averaging (BMA) to obtain a single model with connectivity parameters corresponding to the average connectivity of all models, with each parameter weighted by its model evidence (models with higher evidence contributed more than models with lower evidence).

###### Inference on parameters

After model selection (with BMA), we made inference on the parameters to assess: i) significance of the driving visual input (one-sample *t*-test), ii) significance of endogenous connectivity between the bm-pSTS and the EBA (one-sample *t*-test), iii) significance of modulatory parameters associated with facing dyads and with non-facing dyads (one-sample *t*-test), and iv) differences in the modulation of endogenous connectivity between facing and non-facing dyads (pairwise *t*-test) (Stephan et al. 2009, 2010; Seghier 2010). Finally, independent-sample *t*-tests were used to test significant differences in modulatory parameters between connection links (EBA→bm-pSTS *vs*. bm-pSTS→EBA).

### Study 2: Match-to-sample task

#### Participants

Twenty healthy participants (12 females, mean age 25 ± 5 years) external to the fMRI study, with normal or corrected-to-normal vision were involved in Study 2 as paid volunteers. All signed an informed consent form approved by the local ethics committee.

#### Stimuli

Study 2 included the point-light displays of facing and non-facing dyads used in the fMRI study (10 facing dyads and 10 non-facing dyads and the horizontally-flipped versions for a total of 40 displays). A new set of displays was created as a control condition, by inverting the above displays upside-down. Thus, the total number of stimuli was 80. Finally, a 2-s masking video was created, consisting of white dots moving horizontally on a black background.

#### Experimental design and procedure of the visual recognition task

The experiment included four runs. Each run included 10 trials for each of the four conditions (upright and inverted, facing and non-facing dyads), presented in random order (total of 160 trials). Each stimulus was presented twice throughout the experiment (80 original stimuli and their flipped version). Each trial consisted of two consecutive presentations of either the same stimulus (same-trial, 50% over a run) or of two different stimuli (different trials). The two stimuli, whether identical or different, always belonged to the same condition; for example, if the first stimulus (i.e., sample) was an upright facing dyad, the following stimulus (i.e., probe) could be the same or a different upright facing dyad. The sample (2 s) was followed by a masking video (2 s) and finally by the probe (2 s). After the probe, a blank remained on the screen until the participant gave a response. The next trial began after 500 ms. Participants were instructed to judge if the probe was identical or different with respect to the sample, by pressing one of two keys on a keyboard in front of them (“q” for “same” and “m” for “different”), as soon as possible. Participants sat on a height-adjustable chair, 60 cm away from a computer screen, with their eyes aligned to the center of the screen (17-in. CRT monitor; 1024 × 768 pixel resolution; 85-Hz refresh rate). Stimuli on the screen did not exceed 7° of visual angle. Between two runs, participants were invited to take a break. A run of familiarization preceded the experiment (20 trials). The experiment lasted approximately 40 min. Stimulus presentation and response collection (accuracy and reaction times, RTs) were controlled with Psychtoolbox (Brainard 1997) through MATLAB.

#### Data analyses

Mean accuracy (mean proportion of correct responses) and RTs were entered in two separate ANOVAs with within-subjects factors: Spatial relation (facing *vs*. non-facing) and Orientation (upright *vs*. inverted). The RT analysis included trials in which the participant’s response was correct and within 2.5 standard deviations from the individual mean. Pairwise comparisons between critical conditions were performed using pairwise *t*-tests (α = 0.05, two-tailed).

## RESULTS

### Study 1: fMRI results

#### Effect of spatial relation

##### Whole-brain analysis

Watching movies depicting two bodies acting and moving toward one another (facing dyads), in contrast to two bodies acting and moving away from one another (non-facing dyads), elicited increased neural activity in a distributed brain network encompassing the lateral occipital cortex, the bilateral middle/superior temporal cortex, extending to the temporoparietal junction, the left dorsolateral prefrontal cortex, the left inferior frontal gyrus (pars opercularis), the bilateral precentral gyrus, and the right insula (see Table 1). The opposite contrast (non-facing *vs*. facing dyads) revealed no cluster.

**Table 1.** Results of the whole-brain contrast facing *vs*. non-facing dyads (*p* < 0.001, uncorrected).

| Hemisphere | Brain regions | MNI coordinates |  |  | *t* value |
| --- | --- | --- | --- | --- | --- |
|  |  | x | y | z |  |
| L | Posterior Middle/Superior Temporal Cortex | -49 | -57 | 11 | 5.63 |
| R | Posterior Middle/Superior Temporal Cortex | 43 | -57 | 9 | 4.02 |
| L | Lateral Occipital Cortex | -47 | -73 | 6 | 6.19 |
| R | Lateral Occipital Cortex | 43 | -64 | 6 | 4 |
| L | Primary Motor Cortex | -53 | -15 | 39 | 4.78 |
| R | Primary Motor Cortex | 47 | -15 | 39 | 3.80 |
| L | Premotor Cortex | -53 | -5 | 42 | 5.26 |
| R | Premotor Cortex | 50 | -5 | 35 | 4.53 |
| L | Temporoparietal Junction | -51 | -68 | 9 | 6.45 |
| L | Inferior Frontal Gyrus, pars opercularis | -53 | 8 | 17 | 4.66 |
| L | Dorsolateral Prefrontal Cortex | -40 | 1 | 33 | 4 |
| R | Insula | 43 | -15 | 10 | 4.81 |

*Note.* MNI = Montreal Neurological Institute, L = left, R = right hemispheres.

##### ROI analysis

Whole-brain analysis highlighted stronger activity for facing *vs*. non-facing dyads in a widespread network encompassing the bilateral pSTS and the lateral occipital cortex. To characterize the functional correspondence of the effect of spatial relation with visual processing of body and body motion, we examined the activations in the functionally defined bm-pSTS and EBA. The PPA and the EVC were also examined, to assess the specificity of effects found in the target ROIs. Results showed that the spatial relation between bodies in a dyad affected the response in the bm-pSTS and EBA, but not in the PPA and EVC (Figure 3A). Confirming this observation, the 2 Hemisphere × 4 ROI × 2 Spatial relation ANOVA revealed the main effects of spatial relation, *F*(1,38) = 31.06, *p* < 10^−5^, η_p_^2^ = 0.46, and ROI, *F*(3,114) = 116.99, *p* < 10^−33^, η_p_^2^ = 0.76, but no effect of hemisphere, *F*(1,38) = 0.005, *p* = 0.95, η_p_^2^ = 0.0002. Also significant were the interactions between ROI and hemisphere, *F*(3,114) = 5.42, *p* = 0.002, η_p_^2^ = 0.13, and ROI and spatial relation, *F*(3,114) = 35.24, *p* < 10^−15^, η_p_^2^ = 0.49; the interaction between spatial relation and hemisphere did not reach significance, *F*(1,38) = 3.25, *p* = 0.08, η_p_^2^ = 0.08. All significant main effects and interactions were qualified by the three-way interaction, *F*(3,114) = 9.87, *p* < 10^−5^, η_p_^2^ = 0.21, implying that the effect of spatial relation varied across ROIs and hemispheres. Pairwise *t*-tests showed a difference in the neural response to facing *vs*. non-facing dyads, such that the activity for facing dyads was stronger than for non-facing dyads, in the bilateral bm-pSTS (left: *t*(19) = 5.15, *p* < 0.0001; right: *t*(19) = 4.14, *p* = 0.0006) and EBA (left: *t*(19) = 5.47, *p* < 0.0001; right: *t*(19) = 3.75, *p* = 0.001) but not in the PPA (left: *t*(19) < 1, *ns;* right: *t*(19) < 1, *ns*) and the EVC (left: *t*(19) = 1.69, *p* = 0.11; right: *t*(19) = 1.87, *p* = 0.08).

**Figure 3.**
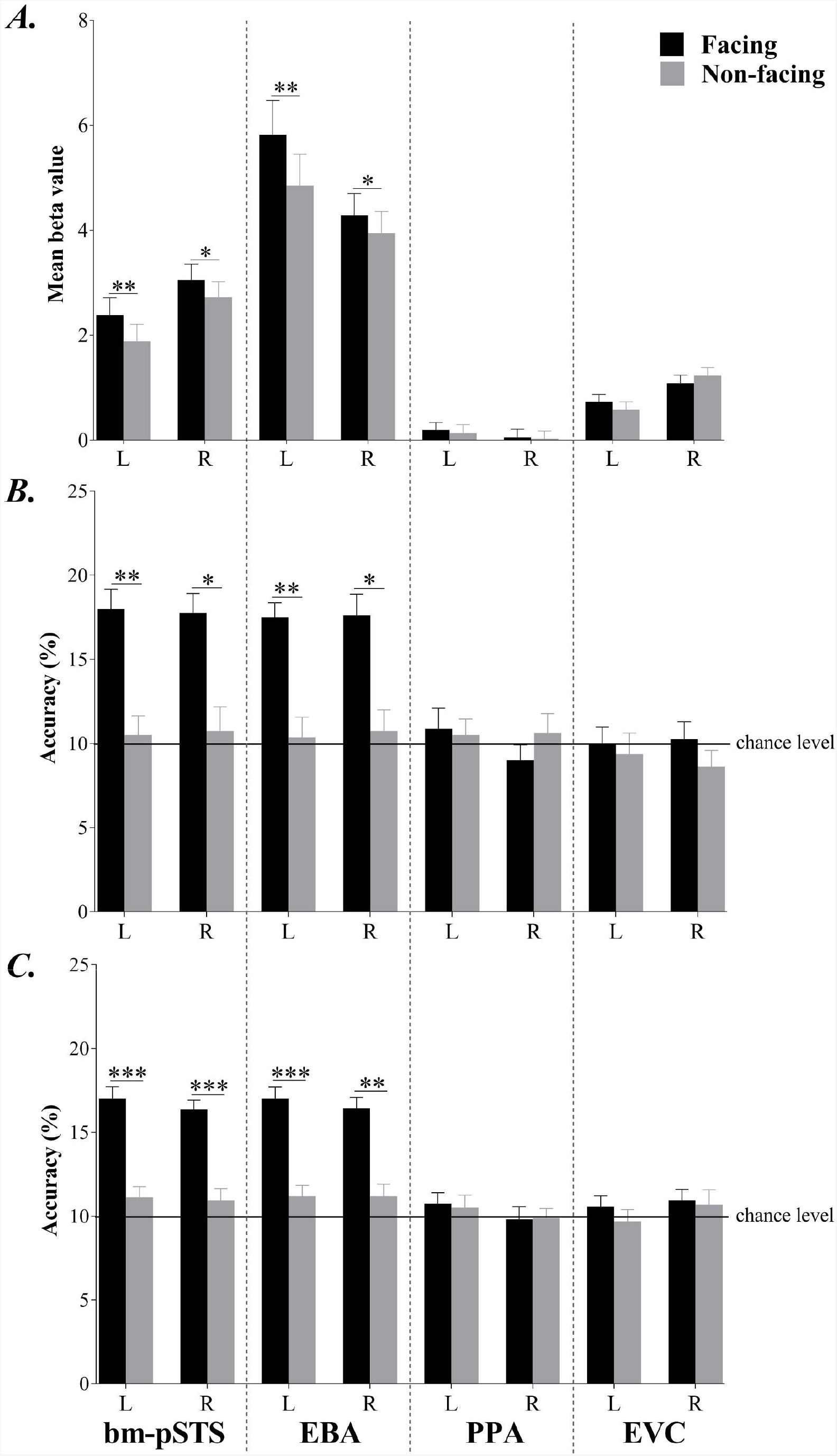
A. Mean beta values (± within-subjects normalized standard error of the mean, SEM) across participants, in each individually defined ROI (bm-pSTS, EBA, PPA and EVC), in response to facing *vs*. non-facing dyads. B. Mean accuracy of classification (mean ± SEM) of facing and non-facing dyads in the ROIs. C. Mean accuracy of classification (mean ± SEM) of single bodies in facing and non-facing dyads in the ROIs. The horizontal line represents the chance level (10%). Asterisks indicate above chance classification: * *p* < 0.001, ** *p* < 10^−4^, *** *p* < 10^−6^ (α = 0.05, two-tailed).

In summary, the whole-brain and the ROI analyses highlighted stronger response to dyads with bodies moving toward (*vs*. away from) each other, in a network of regions that encompassed the bilateral pSTS and EBA, overlapping with the functional areas specialized to visual perception of bodies (EBA) and body motion (bm-pSTS). In the following, we examined the neural representation of dyads as wholes, and of single bodies in the dyads, as a function of the spatial relation (facing *vs*. non-facing) between the two bodies, using MVPA.

#### Representation of facing vs. non-facing dyads in the ROIs

We tested whether the increased activation for facing *vs*. non-facing dyads in the bm-pSTS and EBA was accompanied by a better (i.e., more accurate) representation of the former category of stimuli. Using MVPA, we tested, in each ROI, how well a neural pattern representing a given dyad could be discriminated from the others of the same category (facing or non-facing dyads). We found that in the bm-pSTS classification accuracy was above chance for facing dyads (*bm-pSTS* left: mean accuracy 18% ± 1.2 SEM, *t*(19) = 7.01, *p* < 10^−7^, ES = 1.6; right: 17.7% ± 1.2, *t*(19) = 6.85, *p* < 10^−7^, ES = 1.5) but not for non-facing dyads (*bm-pSTS* left: 10.5% ± 1.1, t(19) < 1, *ns*; right: 10.7% ± 1.4, *t*(19) < 1, *ns*) (Figure 3B). Similar results were found in the EBA, with classification accuracy significantly above chance for facing dyads (left: 17.5% ± 0.8, *t*(19) = 9.02, *p* < 10^−10^, ES = 2.02; right: 17.6% ± 1.2, *t*(19) = 6.21, *p* < 10^−6^, ES = 1.38) but not for non-facing dyads (left: 10.4% ± 1.2, *t*(19) < 1, *ns*; right: 10.7% ± 1.3, t(19) < 1, *ns*). Moreover, in the bilateral bm-pSTS and EBA, classification accuracy was significantly higher for facing dyads than for non-facing dyads (*bm-pSTS* left: *t*(19) = 4.71, *p* < 0.0001, ES = 1.48; right*: t*(19) = 3.92, *p* < 0.001, ES = 1.22; *EBA* left: t(19) = 4.98, *p* < 0.0001, ES = 1.53; right: *t*(19) = 3.9, *p* < 0.001, ES = 1.25). The same analysis in the PPA and EVC yielded no significant effects (*PPA*, facing dyads – left: 10.8% ± 1.2, *t*(19) < 1, *ns*; right: 9% ± 0.9, *t*(19) = 1.12, *p* = 0.27; non-facing dyads – left: 10.5% ± 0.9, *t*(19) < 1, *ns*; right: 10.6% ± 1.1, *t*(19) < 1, *ns*; *EVC*, facing dyads - left: 10% ± 1, *t*(19) < 1, *ns*; right: 10.2% ± 1.1, *t*(19) < 1, *ns*; non-facing dyads - left: 9.4% ± 1.2, *t*(19) < 1, *ns*; right: 8.6% ± 1, *t*(19) = 1.41, *p* = 0.16).

In summary, relative to a non-facing configuration, the face-to-face positioning of bodies and bodily movements enhanced the representation of the whole visual scene, selectively in the EBA and bm-pSTS.

#### Representation of individuals in dyads

Using MVPA with a multi-class cross-decoding scheme, we measured, in each ROI, how well single bodies, presented to the classifier during training, could be discriminated in the (facing or non-facing) dyads presented during the test. Results showed that single bodies were discriminated accurately from patterns of facing dyads, and only in the bm-pSTS and EBA (Figure 3C). In the bm-pSTS, classification accuracy was significantly above chance for the test on facing dyads (*bm-pSTS* left: 17% ± 0.7, *t*(19) = 9.84, *p* < 10^−11^, ES = 2.20; right: 16.4% ± 0.5, *t*(19) = 12.18, *p* < 10^−13^, ES = 2.72), while it was not significantly different from chance for the test on non-facing dyads (*bm-pSTS* left: 11.1% ± 0.6, *t*(19) = 1.83, *p* = 0.081; right: 10.9% ± 0.7, *t*(19) = 1.35, *p =* 0.184). A similar pattern of results was found in the EBA, with classification accuracy significantly above chance for the test on facing dyads (left: 17% ± 0.7, *t*(19) = 10.03, *p* < 10^−11^, ES = 2.24; right: 16.4% ± 0.6, *t*(19) = 10.08, *p* < 10^−11^, ES = 2.26), but not for the test on non-facing dyads (left: 11.2% ± 0.6, *t*(19) = 1.91, *p* = 0.06; right: 11.1% ± 0.7, *t*(19) = 1.56, *p* = 0.13). Accuracy was significantly higher for classification of facing-dyad patterns *vs*. non-facing dyad patterns in both the bm-pSTS (left: *t*(19) = 6.27, *p* < 10^−6^, ES = 1.97; right: *t*(19) = 6.29, *p* < 10^−6^, ES = 1.96) and the EBA (left: *t*(19) = 6.21, *p* < 10^−6^, ES = 1.95; right: *t*(19) = 5.53, *p* < 10^−5^, ES = 1.74). The same analysis in the PPA and EVC yielded no significant effects (*PPA*, facing dyads – left: 10.7% ± 0.6, *t*(19) = 1.20, *p* = 0.24; right: 9.8% ± 0.7, *t*(19) < 1, *ns*; non-facing dyads – left: 10.5% ± 0.7, *t*(19) < 1, *ns*; right: 9.9% ± 0.6, *t*(19) < 1, *ns*; *EVC*, facing dyads – left: 10.6% ± 0.6, *t*(19) < 1, *ns*; right: 10.9% ± 0.7, *t*(19) = 1.45, *p* = 0.15; non-facing dyads – left: 9.7% ± 0.7, *t*(19) < 1, *ns*; right: 10.7% ± 0.9, *t*(19) < 1, *ns*).

In summary, selectively in the EBA and bm-pSTS, single bodies seen in face-to-face configuration was represented better than the very same bodies seen in a non-facing (back-to-back) configuration.

#### Modulation of effective connectivity between the EBA and bm-pSTS by spatial relations

DCM analysis was performed to address how visual perception of facing dyads and non-facing dyads modulated the connectivity between the EBA and the bm-pSTS underlying perception of dynamic single bodies. In the defined model space (see Figure 2), models 2 and 3, which involve bidirectional connections between the EBA and bm-pSTS, with modulation by the experimental context on one (EBA→bm-pSTS, model 2) or both connections (EBA→bm-pSTS and bm-pSTS→EBA, model 3), outperformed other models in accounting for the data in the non-facing condition (xp = 0.47) and in the facing condition (xp = 0.46), respectively (Figure 4A-B). However, since neither model had a xp above 0.9, BMA was applied over the model spaces (i.e. “facing dyads” and “non-facing dyads” model spaces) to obtain, for each space, a single model with weighted average connectivity and modulatory parameters.

**Figure 4.**
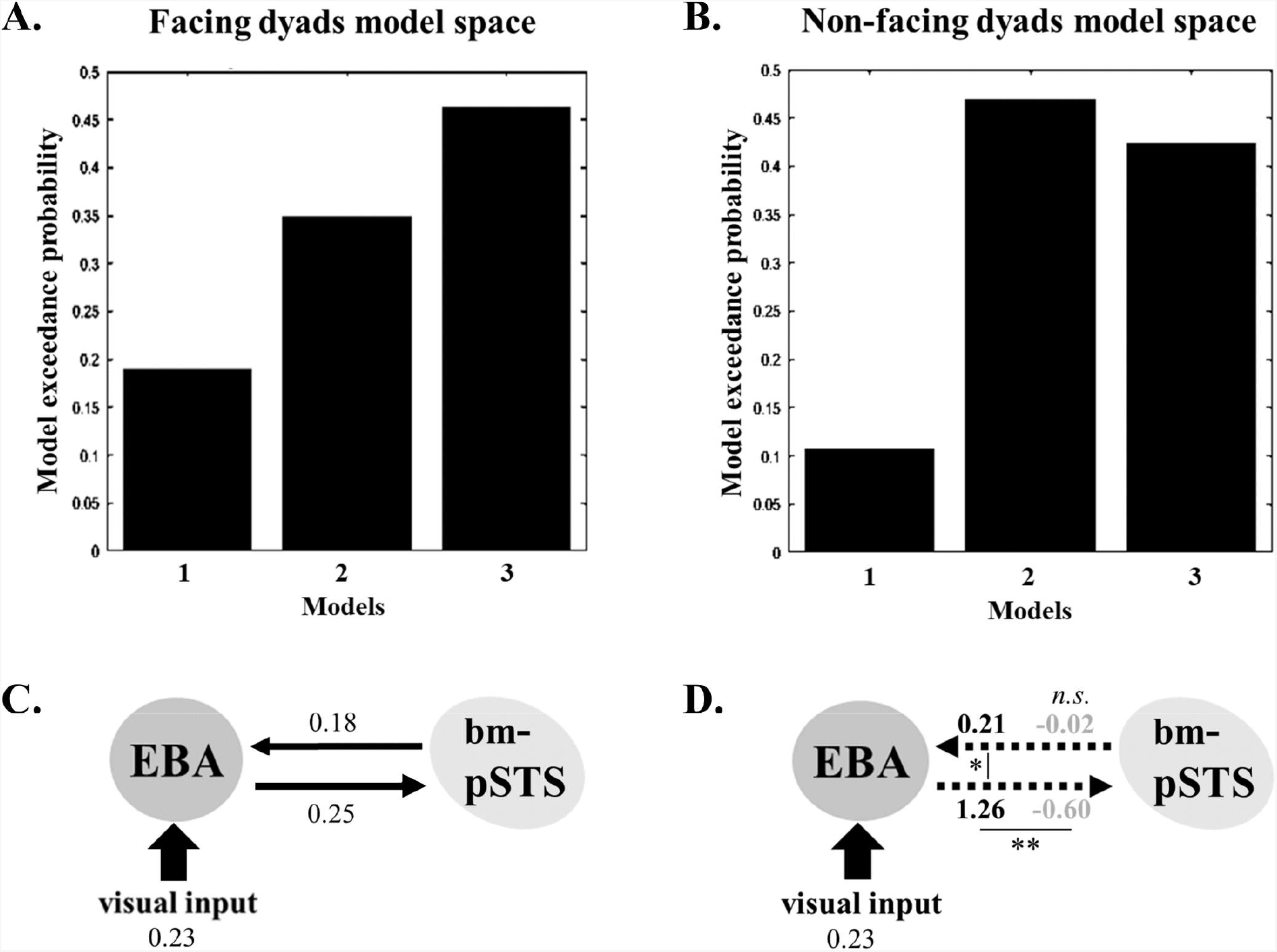
A-B. Exceedance probabilities (xp) of the three models tested for perception of facing dyads (A) and non-facing dyads (B). C. Bayesian Model Averaging (BMA) parameters analysis results for endogenous connections (expressed in Hz). D. BMA parameters analysis results for modulatory connections (expressed in Hz). In bold black, modulation parameters for facing dyad perception and in bold grey, modulation parameters for non-facing dyad perception (i.e., how much endogenous connections are influenced by facing/non-facing dyads perception). Solid thickest black arrows represent driving visual input (single bodies). Black arrows represent endogenous connectivity and dotted arrows represent endogenous connections modulated by perception of facing or non-facing dyads. * *p* < 0.001, ** *p* < 10^−5^ (α = 0.05, two-tailed). *n.s*. = parameters not significant.

We first addressed the significance of driving input parameter and endogenous connection parameters against the null hypothesis (i.e., parameters equal to zero), using one-sample *t*-tests. Results showed significant driving effect of the visual input (single bodies) in the EBA (*t*(19) = 7.91, *p* < 10^−8^, ES = 1.75), and significant increase in the endogenous coupling between the EBA and the bm-pSTS (EBA→bm-pSTS: *t*(19) = 3.99, *p* < 0.001, ES = 0.89; bm-pSTS→EBA: *t*(19) = 3.22, *p* = 0.003, ES = 0.72). Significant positive connectivity from the EBA to the bm-pSTS (0.25 Hz) and from the bm-pSTS to the EBA (0.18 Hz) indicated that, during body perception, an increase in the EBA activity resulted in an increase in bm-pSTS activity and that, in turn, increased bm-pSTS activity resulted in increased EBA activity (Figure 4C, Table 2).

**Table 2.** Descriptive statistical values (mean and standard deviation, SD) for the different parameters, i.e. driving visual input, endogenous connections and modulations, and the statistical values for the *t*-tests, including the *t*-values, *p*-values and effect-sizes (ES). Bold values represent significant values (significantly different from zero).

| Parameters | mean | SD | *t* | *p* | ES |
| --- | --- | --- | --- | --- | --- |
| **Driving visual input** |  |  |  |  |  |
| Single bodies | **0.23** | **0.13** | **7.91** | $p < 10^{-8}$ | **1.75** |
| **Endogenous connections** |  |  |  |  |  |
| EBA→bm-pSTS | **0.25** | **0.28** | **3.99** | $p < 0.001$ | **0.89** |
| bm-pSTS→EBA | **0.18** | **0.25** | **3.22** | $p = 0.003$ | **0.72** |
| **Modulation by Facing dyads** |  |  |  |  |  |
| EBA→bm-pSTS | **1.26** | **1.14** | **4.94** | $p < 0.0001$ | **1.10** |
| bm-pSTS→EBA | **0.21** | **0.14** | **6.71** | $p < 10^{-7}$ | **1.51** |
| **Modulation by Non-facing dyads** |  |  |  |  |  |
| EBA→bm-pSTS | **-0.60** | **1.10** | **2.44** | $p = 0.021$ | **0.54** |
| bm-pSTS→EBA | -0.02 | 0.14 | 0.79 | $p = 0.44$ | 0.18 |

We then evaluated the modulatory effect of perceiving facing dyads and non-facing dyads, on the connection strength between the EBA and the bm-pSTS, with one-sample *t*-tests against the null hypothesis (i.e., parameters are equal to zero). Modulation by facing dyads was significant for both connections (EBA→bm-pSTS: *t*(19) = 4.94 *p* < 0.0001, ES = 1.10; bm-pSTS→EBA: *t*(19) = 6.71, *p* < 10^−7^, ES = 1.51). Positive modulation of connectivity from the EBA to the bm-pSTS (1.26 Hz) and from the bm-pSTS to the EBA (0.21 Hz) suggests that perception of facing dyads increased the connection strength between the EBA and the bm-pSTS. A *t*-test showed that modulatory effect of facing dyads was stronger for the connection from the EBA to the bm-pSTS than for the connection from the bm-pSTS to the EBA (*t*(19) = 4.09, *p* < 0.001, ES = 1.29).

Conversely, modulation by non-facing dyads was only significant for the connection from the EBA to the bm-pSTS, (*t*(19) = 2.44, *p* = 0.021, ES = 0.54). This modulation was negative (−0.60 Hz), suggesting that non-facing dyads perception decreased the connection strength from the EBA to the bm-pSTS (Figure 4D and Table 2). Pairwise comparisons between modulatory parameters for facing and non-facing dyad revealed that facing dyads (*vs*. non-facing) had a stronger modulatory effect on the endogenous connectivity from the EBA to the bm-pSTS (mean difference ± SD difference: 1.86 ± 0.04, *t*(19) = 5.25, *p* < 10^−5^, ES = 1.66). As modulatory parameters add up to endogenous parameters, these results indicate that, the connectivity from the EBA to the bm-pSTS was greater in the facing-dyad, than in the non-facing dyad condition. We could not compare facing and non-facing dyad modulatory parameters for the opposite connection (bm-pSTS→EBA) because non-facing dyads had no significant (modulatory) effect on that connection. However, the significant positive modulation of this connection by facing dyads suggests that the connection strength from the bm-pSTS to the EBA was greater in the facing-dyad, than in the non-facing dyad condition.

### Study 2: Visual recognition of facing *vs*. non-facing dyads

fMRI results showed a better representation of facing *vs*. non-facing dyads in the EBA and the bm-pSTS. Results of the match-to-sample task captured the same advantage in the participants’ behaviour. Confirming a tuning of human vision for facing dyads, the 2 Spatial relation (facing *vs*. non-facing) × 2 Orientation (upright *vs*. inverted) repeated-measures ANOVA on accuracy values showed no significant effects of spatial relation, *F*(1,38) = 2.63, *p* = 0.112, η_p_^2^ = 0.06, and orientation, *F*(1,38) = 0.04, *p* = 0.835, η_p_^2^ = 0.001, but a significant interaction between the two, *F*(1,38) = 20.72, *p* < 0.0001, η_p_^2^ = 0.34. The interaction reflected higher accuracy for facing *vs*. non-facing dyads when presented upright, *t*(19) = 5.93, *p* < 10^−5^, ES = 1.31, but not difference between the two conditions with stimuli presented upside-down, *t*(19) = 1.24, *p* = 0.231 (see Figure 5). This pattern of results maintains that the performance difference between facing and non-facing dyad conditions was due to perception of different configurations of bodies/body movements, as opposed to low- or mid-level visual features, which were preserved in inverted stimuli. The same ANOVA on RTs revealed an effect of orientation, *F*(1,38) = 28.80, *p* < 10^−5^, η_p_^2^ = 0.42 (i.e., participants were generally slower in recognizing inverted *vs*. upright stimuli), but no effect of spatial relation, *F*(1,38) = 0.0006, *p* = 0.979, or interaction, *F*(1,38) = 0.02, *p* = 0.872.

**Figure 5.**
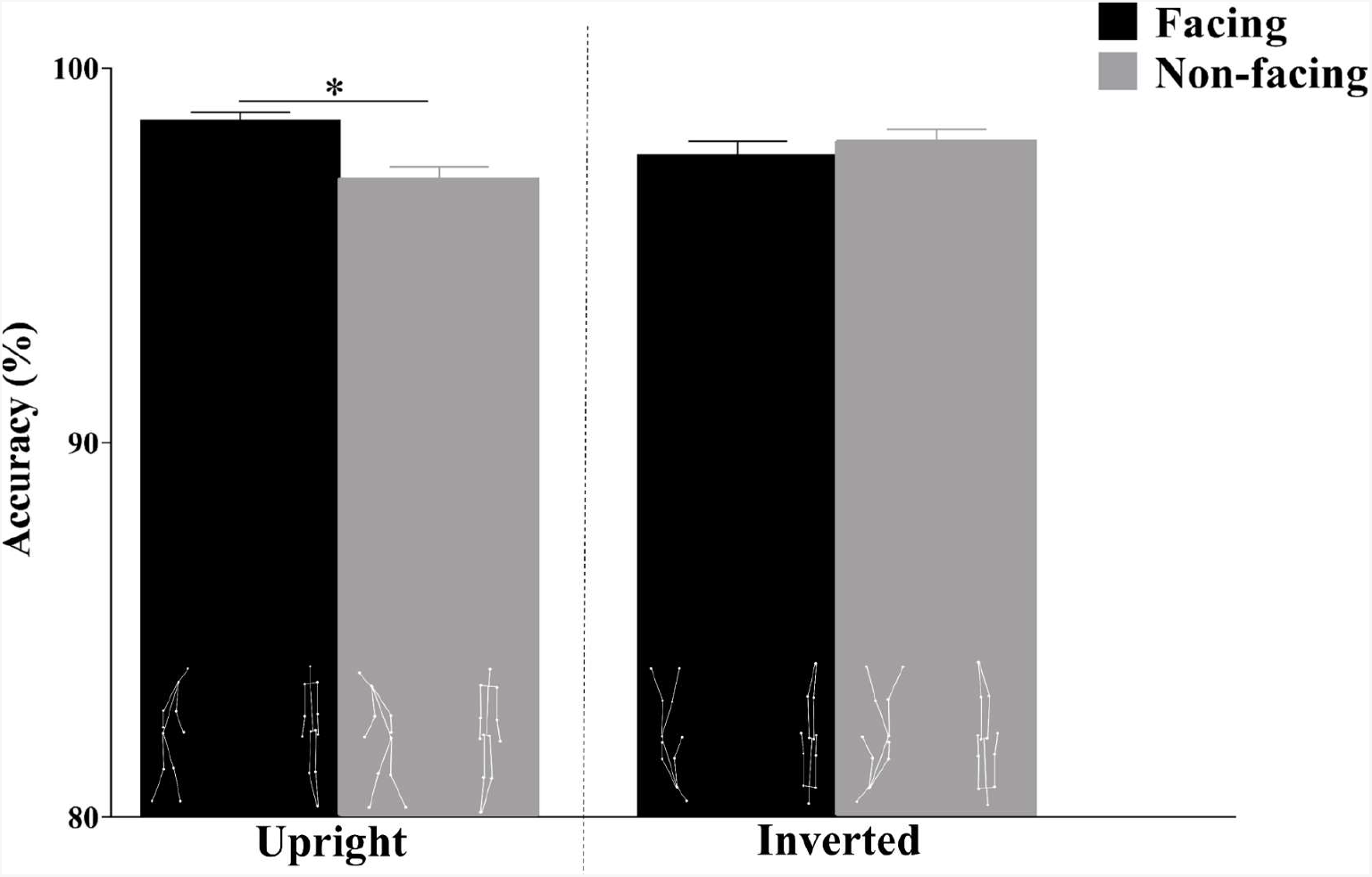
Accuracy (mean ± SEM) of recognition of upright and inverted facing *vs*. non-facing dyads. * *p* < 10^−5^ (α = 0.05, two-tailed).

## DISCUSSION

### Brain areas for body and body-motion perception are tuned to facing body dyads

We asked whether visual areas dedicated to body form and body motion perception can hold the representation of multiple moving bodies and capture relations between them. Since these tasks are fundamental to construct a representation of events such as social interactions, our investigation ultimately addressed how visual perception areas contribute to event formation.

Previous research had shown that, in addition to their primary functions (i.e., representation of body form in the EBA and of body motion in the bm-pSTS), the EBA encodes the orientation of single bodies (leftward *vs*. rightward) and the bm-pSTS encodes the direction of body motion (e.g., moving forward *vs*. backward) (Vangeneugden et al. 2014). The current study shows that the very same brain areas compute information about *relative* body orientation and motion direction, that is, a body’s orientation and motion direction with respect to another body. Indeed, we found that spatial relations between two bodies affected the representation of body and bodily movements in the EBA and the bm-pSTS, and the neural coupling between them. In particular, perception of bodies acting and moving toward (*vs*. away from) each other increased the fMRI response, and the neural integration of information about body form and motion, which is fundamental for action representation (Grossman and Blake 2002; Gu et al. 2020). The stronger neural response to facing (*vs*. non-facing) dyads in the EBA and bm-pSTS was accompanied by more accurate neural discrimination of the overall scenes featuring facing dyads, and of single body postures/movements in those scenes. Finally, an independent behavioural study (Study 2) related the enhanced representation of facing bodies to particularly high performance in visual recognition of facing (*vs*. non-facing) dyads.

Increased activity for facing dyads in the high-level visual areas and the participants’ performance in the visual recognition task are in keeping with effects previously reported for static images depicting multiple-person scenes. In particular, it has been shown that images of face-to-face bodies increase the activity in the face- and body-selective visual cortex including the EBA and the fusiform areas for face and body perception (Abassi and Papeo 2020). Moreover, behavioural studies targeting performance-based measures of perception, have shown that, relative to other configurations, face-to-face bodies have privileged access to visual awareness in low-visibility conditions (i.e., visual categorization of stimuli with fast presentation and backward masking; Papeo et al. 2017a), and are more likely to recruit attention in visual search though a crowd (Papeo et al. 2019; Vestner et al. 2019).

Those previous results have suggested a particular visual sensitivity, or tuning, to multiple-body configurations with visuo-spatial relations that are reliably associated with social interaction, such as *facingness*, spatial proximity, and approaching signals (see Papeo 2020 for discussion). The new results presented here generalize to dynamic multiple-person scenarios and to the biological motion-perception cortex, the visual specialization for facing-body configurations. In doing so, they suggest that such specialization is not just a functional property of brain areas for person perception, but it is a property of the larger network that contributes to the construction of action representation.

### Neural interactions during visual perception of bodies moving toward vs. away from each other

Analogous response profiles in the EBA and bm-pSTS (but not in other visual areas such as the PPA and the EVC) and the estimates of effective connectivity indicate that computations in the two areas are integrated in a processing that is especially driven by spatial cues of interaction in multiple-body scenarios.

Visual mechanisms for perception of body form and body motion mutually contribute to constructing the representation of bodily actions (Grossman and Blake 2002; Gu et al. 2020), an interplay that is particularly emphasized during perception of point-light displays, where motion defines the body shape (Johansson 1973; Mather et al. 1992; Neri et al. 1998). The results reported here confirmed the basic coupling between the EBA and the bm-pSTS during human movement perception (Gu et al. 2020). In addition, the results showed that the (bidirectional) functional coupling between body-form and body-motion processing areas increased for multiple face-to-face bodies, suggesting strong engagement of the action processing network. On the contrary, for non-facing bodies, the connectivity strength between the EBA and the pSTS decreased, as if the scene became less relevant for action-related processing (and perhaps more relevant for other types of analyses).

The current results suggest that the network described for human movement perception houses a mechanism for processing multiple-person scenarios, more akin to what the perceptual system must parse in real-world scenes. This mechanism relies on the encoding of spatial relations not just between parts of a single body to define posture and action, but also between different bodies in the scene, possibly to define interaction.

### Neural sharpening of facing dyads and bodies

Discrimination of scenes involving non-facing dyads and of single bodies in non-facing dyads was at chance in both the EBA and the bm-pSTS. This implies that holding the representation of two bodies moving and acting simultaneously, during color-change detection, which diverted the participants’ attention away from the bodies/actions, is not an easy task. How did spatial relations in facing dyads enhance body representation?

Based on participants performance in visual search, it has been proposed that face-to-face bodies form a *hot spot* that strongly recruits attention (Vestner et al. 2020); enhanced visual representation could be a consequence of such attentional advantage. On another -not mutually exclusive-account, a configuration of two face-to-face bodies could match the observer’s internal template, or expectation about the spatial arrangement of nearby bodies. In this perspective, better discrimination of facing (*vs*. non-facing) bodies is reminiscent of the neural enhancement, or *sharpening*, reported in the visual cortex, for a single object seen in a meaningful or expected visual context, such as a car on a road (*vs*. a car in isolation; Brandman and Peelen 2017; see also Abassi and Papeo 2020; Heilbron et al. 2020). Effects of sharpening are the neural counterpart of visual context effects documented in classic studies on visual object recognition (e.g., word superiority effect; Reicher 1969; face superiority effect; Homa et al. 1976), and more recently extended to perception of social interactions (Neri et al. 2006). In this line of research, it has been shown, for instance, that a single letter is encoded better when seen in a familiar word than in a unknown word (Reicher 1969), and a nose is encoded better in a face than in a scrambled face (Homa et al. 1976). Likewise, an individual’s action (Neri et al. 2006; Ding et al. 2017) or an individual’s role in the action (agent or patient; Hafri et al. 2013) is recognized more accurately when the individual interacts with another, than when she does not. Enhancing effects of social interaction on body action representation have been shown in the context of meaningful interaction (Neri et al. 2006; Hafri et al. 2013; Ding et al. 2017).

In the current study, bodies faced one another without giving rise to any meaningful interaction. Thus, interpreted in the framework of visual context effects, our results suggest that mere spatial relations associated with interaction, even without a semantically specified interaction, can provide a visual context that enhances and enriches body action representation.

### Facingness versus interaction

Unlike previous research that used meaningful interactions with agents performing complementary or coordinated actions (e.g., Sinke et al. 2010; Centelles et al. 2011; Van den Stock et al. 2015; Isik et al. 2017; Walbrin et al. 2018; Walbrin and Koldewyn 2019), we included stimuli with no temporal contingency and no semantic relatedness between the two body movements. As a result, neither facing nor non-facing dyads displayed meaningful interactions, as also evaluated in the preliminary study for stimulus selection. Thus, a difference between meaningful interaction *vs*. no (or meaningless) interaction can hardly account for the difference in the neural activation for facing *vs*. non-facing dyads, or for better discrimination across different exemplars of facing (*vs*. non-facing) dyads in the EBA and the bm-pSTS. We propose that different representation of facing and non-facing dyads in those visual areas rather captures information available, in a bottom-up fashion, in the physical structure of the stimuli; particularly, the difference between facing and non-facing dyads would reflect the appearance (*vs*. absence) of relational properties that cue interaction, rather than the access to the representation of a semantically specified interaction.

Since we aimed at addressing the visual processing of multiple-body actions, our analysis focused on the visual aspect (the EBA and the bm-pSTS) of the larger network that responded to facing, more strongly than to non-facing dyads (see Table 1). The recruitment of regions associated with processing of social stimuli, and social tasks in general (temporo-parietal junction, dorsolateral prefrontal cortex, inferior frontal gyrus, precentral gyrus, and insula), suggests that face-to-face bodies, more than non-facing bodies, might spontaneously trigger some sort of social inference, even in the absence of a semantically specified interaction content. Future research shall address the overlap between the network described here for facing (*vs*. non-facing) dyads and the network activated by the processing of social interaction contents described elsewhere (Centelles et al. 2011; Baldassano et al. 2016; Isik et al. 2017; Walbrin et al. 2018; Walbrin and Koldewyn 2019). Moreover, extending the effective connectivity analysis to other areas responsive to facing (*vs*. non-facing) dyads will help characterize the relationship (and integration) between the processing of relational cues of interaction (i.e., facingness) in the visual EBA and bm-pSTS, and the processing of social interaction in other, non-visual aspects of the network.

## Conclusions

Seeing bodies facing and moving toward each other increased neural activity in a broad brain network including the EBA and the bm-pSTS, enhanced the representation of individual body postures and movements in those regions, and increased discrimination of the overall visual scene. Different neural response to facing and non-facing stimuli implies that the very same areas for body and body motion perception, also register the presence of multiple bodies in a scene (see Cracco et al. 2019, for a related discussion), and the (spatial) relationship between them, and use that information to enhance the representation of configurations with ‒seemingly interacting– bodies. The above effects occurred together with an increase in the coupling between the EBA and the bm-pSTS during movement perception of facing dyads. Thus, visuo-spatial cues of interaction not only affect the perceptual representation of body form and movements in the dedicated brain areas, but also promote the functional integration between the two systems, a mechanism that is fundamental for action representation.

The integrated EBA/bm-pSTS network for multiple-body movement perception characterized here, could be the gateway to a wider network specialized in the representation of social interaction (Isik et al. 2017; Walbrin et al. 2018; Tarhan and Konkle 2020). To the social interaction network, the integrated activity of the EBA and the bm-pSTS (and possibly other visual areas), would deliver the representation of a multiple-body structure that reflects the relational information in the input. In sum, the minimal network of visual areas encompassing the EBA and the bm-pSTS would give rise to the perceptual rudiment of the to-be representation of a social interaction, based on the physical structure of the input, before inferential processes kick in.

## Author Contributions

EB and LP contributed to the conception and design of the study.

EB, LP and EA contributed to acquisition and analysis of data.

EB and LP contributed to drafting the manuscript or preparing the figures.

All authors approved the final version of the manuscript for submission.

## Conflict of interest declaration

The authors declare no competing financial interests.

## Acknowledgments

This work was supported by a European Research Council Starting Grant to L.P. (Grant number: THEMPO-758473)

## Notes

### Competing Interest Statement

The authors have declared no competing interest.

